# Studying RNA–DNA interactome by Red-C identifies noncoding RNAs associated with repressed chromatin compartment and reveals transcription dynamics

**DOI:** 10.1101/859504

**Authors:** Alexey A. Gavrilov, Anastasiya A. Zharikova, Aleksandra A. Galitsyna, Artem V. Luzhin, Natalia M. Rubanova, Arkadiy K. Golov, Nadezhda V. Petrova, Maria D. Logacheva, Omar L. Kantidze, Sergey V. Ulianov, Mikhail D. Magnitov, Andrey A. Mironov, Sergey V. Razin

## Abstract

Non-coding RNAs (ncRNAs) participate in various biological processes, including regulating transcription and sustaining genome 3D organization. Here, we present a method termed Red-C that exploits proximity ligation to identify contacts with the genome for all RNA molecules present in the nucleus. Using Red-C, we uncovered the RNA–DNA interactome of human K562 cells and identified hundreds of ncRNAs enriched in active or repressed chromatin, including previously undescribed RNAs. We found two microRNAs—MIR3648 and MIR3687 transcribed from the rRNA locus—that are associated with inactive chromatin genome wide. These miRNAs favor bulk heterochromatin over Polycomb-repressed chromatin and interact preferentially with late-replicating genomic regions. Analysis of the RNA–DNA interactome also allowed us to trace the kinetics of messenger RNA production. Our data support the model of co-transcriptional intron splicing, but not the hypothesis of the circularization of actively transcribed genes.

## Introduction

The vast majority of the eukaryotic genome is transcribed to produce a broad range of RNAs, including both protein-coding and non-coding RNAs (ncRNAs) (Hangauer et al. 2014). Early studies revealed significant numbers of chromatin-associated RNAs (Holmes et al. 1972; Kanehisa et al. 1972; Bynum and Volkin 1980). The current results demonstrate that chromatin-associated RNA plays an important role in nuclear organization, chromatin folding, and transcription control (Holoch and Moazed 2015; Li and Fu 2019; Nozawa and Gilbert 2019). Long ncRNAs (lncRNAs, > 200 nt) participate in various biological processes, from regulating enzymatic activities to sustaining genome imprinting and nuclear body biogenesis (Quinn and Chang 2016; Sun et al. 2018). Specific lncRNAs coordinate cell differentiation and other processes related to cell fate choice (Flynn and Chang 2014). Overexpression, lack, or mutation of various lncRNA genes underlie many human diseases (Esteller 2011). Still, particular functions are unclear for the majority of individual lncRNAs, and some lncRNAs may be a product of transcription noise and lack function altogether (Struhl 2007). Currently, the functional roles and mechanisms of action have been convincingly disclosed for only a few lncRNAs, such as XIST, HOTAIR, and TERC (Engreitz et al. 2016; Quinn and Chang 2016). LncRNAs may modulate the chromatin structure by binding and targeting activator or repressor complexes to particular genomic loci (Geisler and Coller 2013; Sun et al. 2018). Because they are physically linked to DNA via transcribing RNA Pol II molecules, lncRNAs may fulfill their function immediately following or during transcription without the need for processing or redistribution. Examples of cis-acting lncRNAs include lncRNAs from imprinted loci, dosage compensation lncRNAs, antisense RNAs, and autoregulatory RNAs (reviewed in (Quinn and Chang 2016)).

Along with lncRNAs, short (<200 nt) ncRNAs may also play a role in regulating gene expression at the transcriptional level. Thus, promoter-associated RNAs transcribed in both directions from the promoters of structural genes are likely to contribute to transcription activation (Guil and Esteller 2012). MicroRNA (miRNA), the canonical function of which is to suppress mRNA translation in the cytoplasm, occurs in the nucleus as well, where these miRNAs may pair with other ncRNAs localized in certain genome regions and trigger repression or activation of these regions (Roberts 2014).

A growing body of evidence implicates ncRNAs in spatial genome organization (Engreitz et al. 2016; Li and Fu 2019; Nozawa and Gilbert 2019). Several studies suggest that enhancer RNAs (eRNA) help to juxtapose an enhancer and its target promoter (Li et al. 2016). Interestingly, the CTCF architectural protein, which plays a key role in organizing 3D genomes in mammalian cells, is also capable of binding a broad range of ncRNAs on the genome scale (Saldana-Meyer et al. 2014; Kung et al. 2015). The Firre lncRNA was found to mediate the colocalization of several genomic regions located on different chromosomes (Hacisuleyman et al. 2014). The XIST RNA, which is necessary for establishing dosage compensation in mammals, shapes the 3D structure of the inactive X chromosome (Galupa and Heard 2018).

All of the examples described above are likely only the tip of the iceberg. Diverse functions of ncRNAs are only beginning to be unraveled. Further progress in disclosing the functions of ncRNAs in gene regulation will depend on the availability of the genome-wide spectrum of RNA associations with chromosomes, the RNA–DNA interactome. The problem has been addressed in several recent studies (Li et al. 2017; Sridhar et al. 2017; Bell et al. 2018). The protocols developed in the studies cited above for characterization of the RNA–DNA interactome are based on proximity ligation of RNA to the neighboring DNA fragments. Here, we developed a modified strategy for adaptor-mediated RNA–DNA proximity ligation that allows mapping of the 3’ and 5’ ends of the RNA molecule associated with a given DNA site. Using this method, we uncovered a variety of ncRNAs associating with active and repressed chromatin. We also used RNA–DNA interaction data to study the transcriptional dynamics of protein-coding genes.

## Results

### Development of Red-C

The Red-C (<u>R</u>NA <u>e</u>nds on <u>D</u>NA <u>c</u>apture) experimental procedure for mapping the RNA–DNA interactome is based on adapter-mediated RNA–DNA ligation in fixed nuclei followed by high-throughput sequencing of the chimeric RNA–DNA molecules (Fig. 1A, Fig. S1A). Briefly, DNA-protein-RNA complexes are fixed with formaldehyde in vivo, DNA is fragmented with a restriction enzyme, and the ends are blunted and A-tailed. RNA 3’ ends are ligated to a bridge adapter containing a biotinylated nucleotide followed by ligation of the opposite ends of the bridges with DNA ends in spatial proximity. RNA–DNA chimeras are purified, and excess DNA is cut off using MmeI restriction enzyme, the recognition site of which is incorporated into the bridge. After biotin pull-down, reverse transcription is initiated from the bridge with template switching at the 5’ end of the RNA (SMART technology (Zhu et al. 2001)), allowing for the incorporation of a custom Illumina adapter. Finally, another Illumina adapter is ligated to the DNA ends, and the chimeras are amplified and paired-end sequenced (Fig. 1A, Fig. S1A). Sequencing of one end identifies the 5’ end of the RNA, whereas sequencing of the other end reports the DNA fragment ligated to this RNA, the bridge adaptor sequence, and the 3’ end of the same RNA.

**Fig. 1.**
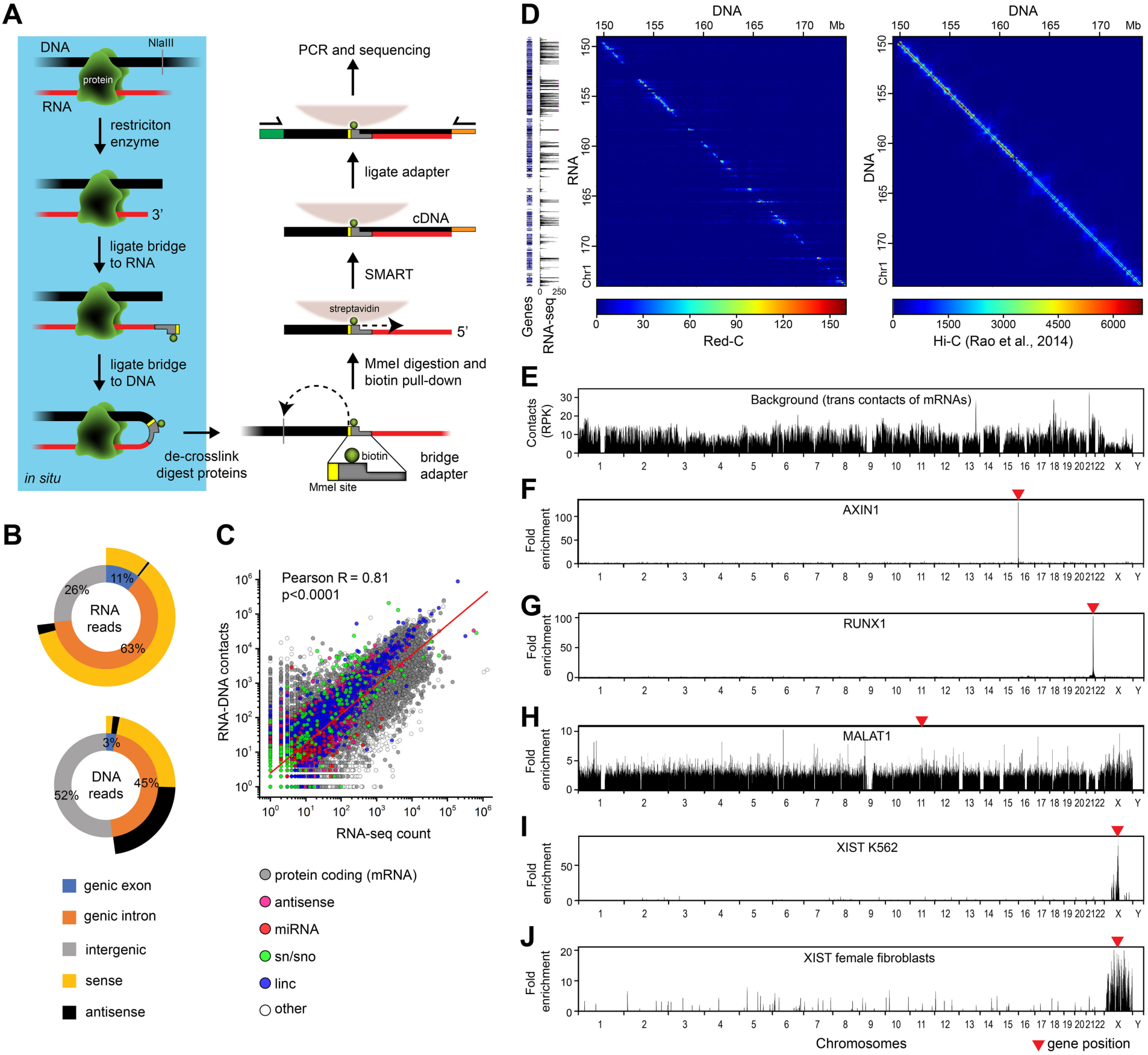
The Red-C technique. (**A**) Outline of Red-C protocol. (**B**) Genomic distribution of DNA and RNA reads extracted from forward and reverse sequencing reads, respectively. As genic, we used RefSeq protein-coding genes that occupy 37% of the genome. Reads having the same direction as the transcript are defined as sense; reads having the opposite direction to the transcript are defined as antisense. (**C**) Correlation of RNA–DNA contacts with RNA-seq signal in K562 cells. Red line, linear regression. (**D**) RNA–DNA (Red-C) and DNA–DNA (K562 Hi-C (Rao et al. 2014)) contact matrices for a region of Chr 1 at a 100 Kb resolution. RNA-seq profile for K562 (1 Kb bins) and gene distribution are shown alongside. (**E**) Background profile in K562 cells. RPK, reads per Kb. (**F–J**) Fold enrichment of selected RNAs compared to the background in K562 cells (**F–I**) and female fibroblasts (**J**). MALAT profile is at 1 Kb resolution; the other profiles are at 100 Kb resolution.

The main difference between Red-C and similar protocols (Li et al. 2017; Sridhar et al. 2017; Bell et al. 2018) is that both the 3’ and 5’ ends of the RNA molecule associated with a given DNA site are identified using SMART, while in the previously published protocols, only the 3’ end is identified. Information about both ends of the ligated RNA chain enables more accurate mapping of RNA and provides more insight into the RNA structure, for example, allowing for the identification of polyadenylated RNAs. The specificity of the Red-C protocol was verified in control experiments with either the omission of the DNA ligation step or treatment of RNA–DNA chimeras with RNase A, resulting in products lacking, correspondingly, DNA or RNA parts (Fig. S1B–E).

We applied the Red-C protocol to uncover the RNA–DNA interactome of the cultured human erythroleukemia cells (line K562). In two biological replicates, we identified 44M unique RNA–DNA contacts (see Table S1 for data processing statistics). Analysis of genomic distribution of RNA and DNA reads showed that the former originated primarily from genic regions and almost exclusively had the same strand orientation as the transcripts, whereas the latter were more uniformly distributed between genic and non-genic regions and, when mapped to genic regions, lacked specificity for the gene strand (Fig. 1B). We then combined the contacts of RNA parts originating from a single gene, thus yielding a whole-genome contact profile for each respective RNA. We also plotted RNA–DNA contact matrices analogous to DNA–DNA contact matrices used in Hi-C analysis (Fig. 1D, Fig. S2). In contrast to Hi-C matrices in which the majority of spatial contacts occur in proximity on the DNA (close to diagonal on the map), the RNA–DNA matrices show a wide distribution of RNA contacts along an extended genomic region (horizontal lines crossing the diagonal). Notably, RNA–DNA contact matrices obtained for individual chromosomes demonstrated good concordance between replicates (Fig. S3, Pearson’s *R* > 0.94). The same is true for contact numbers for individual RNAs (Fig. S4, Pearson’s *R* = 0.96).

The highest number of contacts was observed for mRNAs (31M) and linc and vlinc RNAs (long and very long intergenic non-coding RNAs (St Laurent et al. 2013), 2.7M and 3.2M, respectively). Contacts with the genome were also detected for antisense RNAs, small nuclear and nucleolar (sn and sno) RNAs, miRNAs, piwi RNAs, and other RNA biotypes (Table S2). A considerable number of RNA parts could not be assigned to annotated transcriptional units. Clusters of such RNA parts, which might represent novel RNAs (Table S3, X RNAs), were analyzed along with known RNAs. Notably, the number of captured contacts was proportional to the transcript level, as determined by RNA-seq analysis (Fig. 1C).

To account for differences in chromatin accessibility and ligation efficiency for different genomic sites, we deduced the background level of non-specific ligation based on the distribution of mRNAs in non-parental chromosomes, as suggested by Li et al. (Li et al. 2017) (Fig. 1E). The calculated background level was used for raw signal normalization and fold enrichment calculations (see Methods).

### Distribution of RNAs along the genome

RNAs of different biotypes are characterized by varying distributions of contacts along the genome. For example, the majority of mRNAs are preferentially detected at locations of their synthesis on the chromosome (i.e. in the vicinity of the gene) (Fig. 1F,G), whereas some ncRNAs are also found away from the gene on the same or other chromosomes. In particular, ncRNA MALAT1, which localizes to nuclear speckles and participates in pre-mRNA processing (Tripathi et al. 2010; Sun et al. 2018), is detected in all chromosomes (Fig. 1H). In contrast, ncRNA XIST, which orchestrates X chromosome inactivation in female cells (Galupa and Heard 2018), shows an enrichment over the X chromosome (Fig. 1I). The level of enrichment decreases with an increase in distance from the XIST gene. This phenomenon may be explained by a very high proliferation rate of K562 cells. This cancer cell culture of female origin is nearly void of cells in G0 phase; hence, we may observe the process of XIST expansion over the X chromosome (Engreitz et al. 2013). The possibility of impaired dosage compensation and XIST binding in cancer cells also cannot be ruled out. When we repeated experiments with normal human dermal fibroblasts of female origin, a much more uniform pattern of XIST binding over the entire X chromosome was observed (Fig. 1J). Sporadic signals of XIST on other chromosomes may reflect the probability of these chromosomes being located close to X, though may also be an artifact of the Red-C procedure. The RAP method also detected a small fraction of XIST contacts on autosomes (Engreitz et al. 2013). We also produced a small dataset from male Drosophila S2 cells. Out of 53,378 identified contacts, 1,047 were assigned to roX1 and roX2, ncRNAs that play a vital role in dosage compensation in Drosophila (Samata and Akhtar 2018). Genome binding sites of roX1 and roX2 were extensively studied previously using different approaches including GRID-seq (Li et al. 2017) and ChAR-seq (Bell et al. 2018). Although we had ∼100 times fewer contacts, we were able to reproduce enrichment of roX1 and roX2 in the X chromosome (Fig. S5A) and to obtain binding profiles similar to those generated by GRID-seq and ChAR-seq (Fig. S5B). Taken together, the observations described above confirm the validity of the Red-C protocol because the expected distribution of roX, XIST, MALAT1, and protein-coding RNAs was detected.

For automated analysis of the preferences of individual RNAs for short- and long-range interactions, we assessed the frequency of contacts of each RNA with DNA in several consecutive cis intervals: encoding gene (G); 0–50 Kb upstream and downstream from gene boundaries (S); 50–500 Kb upstream and downstream from gene boundaries (M); 500 Kb–5 Mb upstream and downstream from gene boundaries (L); and > 5 Mb from gene boundaries in the same chromosome (R) (Fig. 2A). We then calculated the ratio of contact frequency in each of the intervals described above (cis contacts) to contact frequency with non-parental chromosomes (trans-contacts, interval T) and presented the ratio as a function of the total number of contacts for each RNA (Fig. 2B). Virtually every RNA showed the highest interaction frequency in the vicinity of the gene and then along the same chromosome (Fig. 2B; see also Fig. 1D, strong signals at the diagonal of the RNA–DNA matrix). However, the degree of enrichment differed drastically for individual RNAs and particular RNA biotypes. For example, sn and sno RNAs demonstrated a low degree of enrichment at the gene-proximal regions and a similar frequency of cis- and trans-contacts (Fig. 2B). T-SNE analysis based on the ratios of contact frequencies in consecutive intervals placed the majority of sn and sno RNAs into a separate cluster (Fig. 2C). XIST also demonstrated a specific behavior, with relatively low preference for gene-proximal areas and higher preference for remote regions of the same chromosome relative to other RNAs, as expected for an RNA distributed over the full length of the parental chromosome (Fig. 2B).

**Fig. 2.**
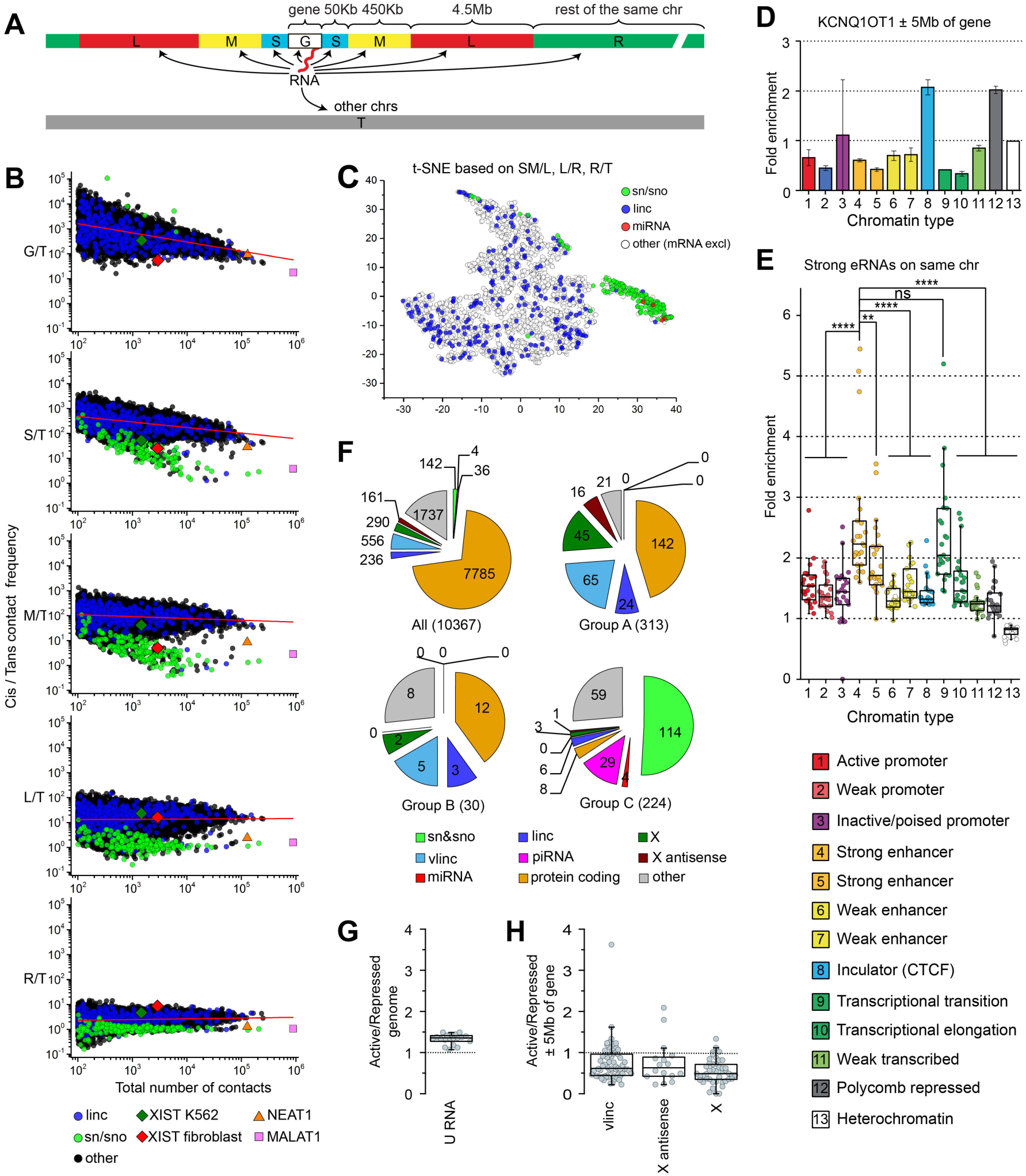
Preferences of RNAs for short- and long-range contacts and different chromatin types in K562 cells. (**A**) Scheme demonstrating analyzed genomic intervals. (**B**) Ratio of contact frequency of individual RNAs with regions of parental chromosome to contact frequency of the same RNAs with the other chromosomes (Y axis) versus total number of contacts (X axis). Graphs from top to bottom show results for different cis intervals as specified in (**A**). RNAs with ≥ 100 contacts are presented. Red line, linear regression. (**C**) T-SNE analysis of RNAs based on ratios between contact frequencies in consecutive intervals. (**D**) Fold enrichment of Kcnq1ot1 at specific chromatin types in the region surrounding Kcnq1ot1 gene (± 5 Mb of gene boundaries) relative to overall contact frequency in this region. Error bars, SEM for two biological replicates. (**E**) Fold enrichment of eRNAs produced from chromatin type 4 and 5 at specific chromatin types within the same chromosome relative to overall contact frequency in the same chromosome. Points represent results for individual chromosomes (*n* = 23, *p*-values are from Tukey’s multiple comparisons test). (**F**) Number of RNAs of a particular biotype in group A (RNAs enriched in gene-proximal areas), group B (XIST-like RNAs), group C (RNAs distributed throughout the genome), and among all analyzed RNAs. (**G,H**) Ratio between contact frequencies in active and repressed chromatin for U RNAs belonging to group C (**G**) and for vlinc, X RNAs, and antisense X RNAs belonging to group A (**H**). Active chromatin is defined as combination of types 1, 2, 4, 5, 6, 7, 9, 10, and 11; repressed chromatin, of types 3, 12, and 13. Contact frequency was determined for the full genome (**G**) or in regions ± 5Mb of gene boundaries (**H**).

### Preferences of RNAs for active and repressed chromatin

We next focused on the preferences of RNA contacts for specific chromatin types. We used the annotation of chromatin states for K562 cells made by Ernst et al. (Ernst et al. 2011). The authors of this study used combinations of chromatin marks to partition the genome into 15 non-overlapping chromatin states typical for active and poised promoters, enhancers, CTCF-dependent insulators, transcribed and Polycomb-repressed regions, *et cetera.* An imprinted antisense RNA Kcnq1ot1 involved in the silencing of several genes in the same locus (Pandey et al. 2008) demonstrated a preference for interaction with Polycomb-repressed regions in the area surrounding the Kcnq1ot1 gene, in agreement with its supposed role in transcriptional repression (Fig. 2D). Additionally, we observed an enrichment of Kcnq1ot1 over CTCF-binding sites (Fig. 2D).

We next focused on enhancer RNAs (eRNAs). Here we define eRNAs as RNAs transcribed from enhancer-specific chromatin states (Table S4). For each chromosomal interval annotated as belonging to a particular chromatin state, we determined the number of contacts established with this interval by eRNAs produced from all over the chromosome, excluding the interval itself. Next, for each chromatin state, we summarized the contacts at all intervals and normalized the sum by the total length of these intervals, thereby obtaining the average contact frequency of eRNAs with particular chromatin states in the parental chromosome. We found that eRNAs produced from strong enhancers showed a preference for other strong enhancers located on the same chromosome (Fig. 2E). This result may reflect the spatial clustering of enhancers. In addition, we observed the enrichment of eRNAs at a transcriptional transition chromatin type (Fig. 2E).

Genomic distribution of an RNA is an important characteristic that may shed light on its potential function. Indeed, RNAs involved in splicing demonstrate a wide spectrum of contacts along all chromosomes, while XIST is spread specifically along the X chromosome. To expand this type of analysis for all RNAs, we developed an algorithm for identification of RNAs with specific genome distribution patterns based on the comparison of contact frequencies between the intervals described above: S+M and L, L and R, and R and T (see Methods and Fig. S6). Using this algorithm, we identified 313 RNAs enriched in gene-proximal areas (group A), 30 XIST-like RNAs enriched over the full length of the parental chromosome (group B), and 224 RNAs distributed along the entire genome (group C) out of 10,367 RNAs with ≥ 500 contacts (Fig. S6, Table S5). Of note, snRNAs, snoRNAs, miRNAs, and piwi RNAs were absent from groups A and B and almost all concentrated in group C (Fig. 2F). Spliceosomal U snRNAs are biased toward active chromatin on a whole-genome scale (Fig. 2G, Fig. S6C). By contrast, vlinc RNAs and antisense X RNAs (newly identified RNAs intersecting a known transcriptional unit and transcribed in the opposite direction) are depleted from group C and significantly overrepresented in group A (Fisher’s exact test *p*-value < 0.0001, Fig. 2F). Remarkably, most of the vlinc RNAs and X RNAs belonging to group A are biased toward repressed chromatin in a 10 Mb region surrounding the gene (Fig. 2H, Fig. S6A).

Of particular interest are MIR3648 and MIR3687 (Fig. 3A). These miRNAs establish contacts genome wide and rank among the first in localization to repressed chromatin and the inactive spatial chromatin compartment annotated previously by eigenvector analysis of Hi-C matrices (Lieberman-Aiden et al. 2009; Rao et al. 2014) (Fig. 3B,C, Fig. S6C, Table S5). Of note, MIR3648 and MIR3687 favor bulk heterochromatin over Polycomb-repressed chromatin (Fig. 3B). They associate with regions of late replication (Fig. 3F,G and data not shown), are depleted from the bodies of transcribed genes, and are enriched in gene deserts (Fig. 3D,E). The frequency of contacts of MIR3648 and MIR3687 with gene-poor chromosome 18 is 2- to 3-fold higher than the genome average (Fig. 3H). Interestingly, MIR3648 and MIR3687 genes are hosted within the 5’ external transcribed spacer of the 45S ribosomal RNA (rRNA) operon (Yoshikawa and Fujii 2016). Based on the length of RNA parts detected in RNA–DNA chimeras, it appears that MIR3648 and MIR3687 act in a form of pre-miRNA rather than processed short miRNA (Fig. 3A). Based on the above observations, it is tempting to speculate that MIR3648 and MIR3687 play a role in heterochromatin formation at the genome scale.

**Fig. 3.**
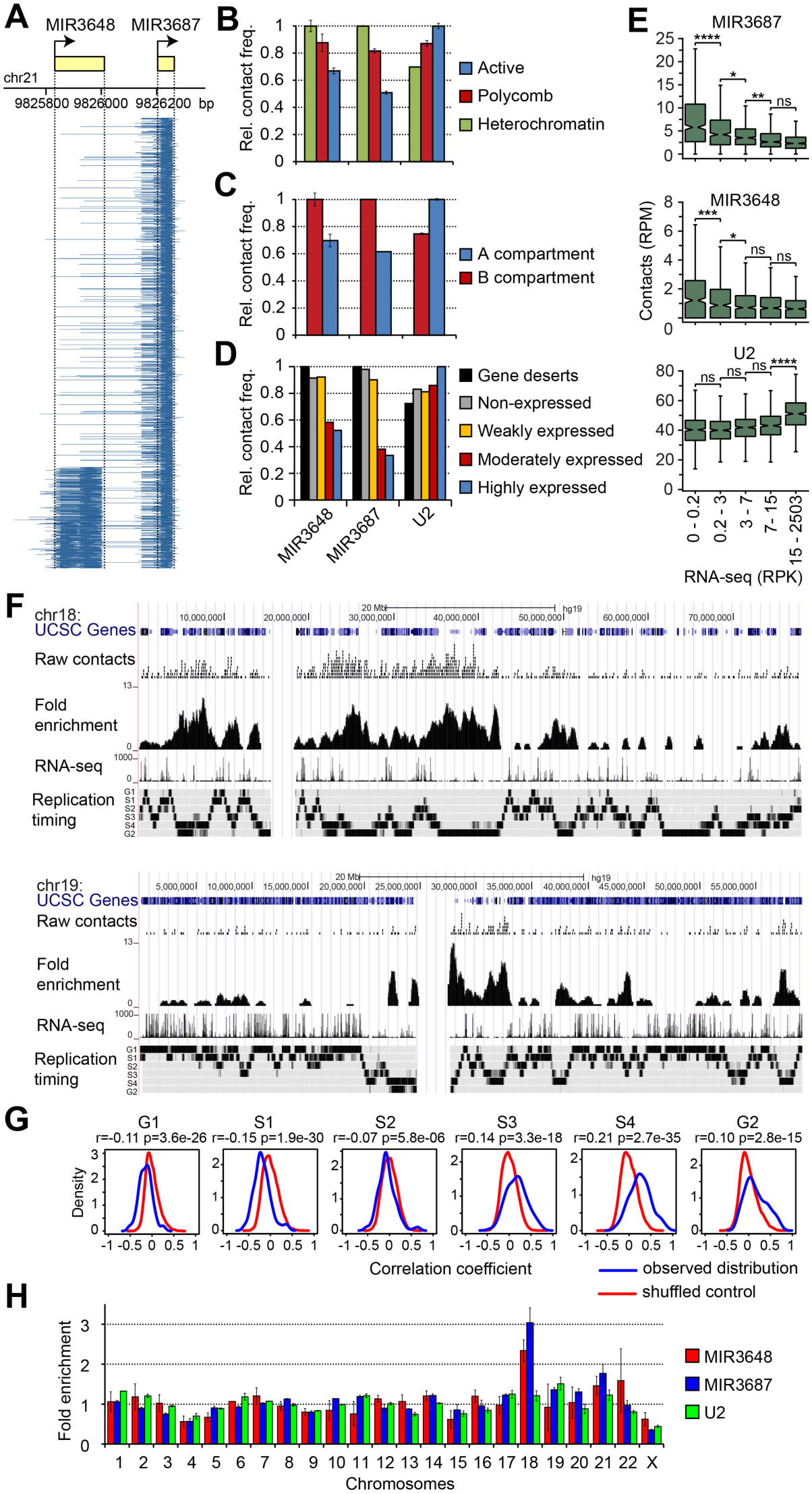
MIR3648 and MIR3687 target inactive chromatin. (**A**) Coverage of the MIR3648/3867 locus by RNA parts of RNA–DNA chimeras. RNA parts are displayed in the stack view in accordance with mapping coordinates of RNA 5’ and 3’ ends. (**B,C**) Frequency of contacts of MIR3648, MIR3687, and U2 with different chromatin types (**B**) and A and B spatial compartments (**C**) determined for the full genome. The maximal contact frequency for a given RNA is taken to be equal to 1. Error bars, SEM for two biological replicates. Active chromatin is defined as combination of types 1, 2, 4, 5, 6, 7, 9, 10, and 11; Polycomb, of types 3 and 12; Heterochromatin, of type 13. A/B compartment track for K562 was obtained from (Rao et al. 2014). (**D**) Frequency of contacts of MIR3648, MIR3687, and U2 with expressed protein-coding genes (divided into three equal groups based on the density of RNA-seq signal), non-expressed protein-coding genes (RNA-seq signal = 0), and gene deserts (regions of > 500 Kb not occupied by any genes). For each RNA, the total number of contacts with genes of each group and gene deserts was determined, normalized by the total length of genes in the group and gene deserts, and presented relative to the maximal value for a given RNA (taken equal to 1). (**E**) Contacts of MIR3648, MIR3687, and U2 with 1 Mb genomic bins divided into five equal groups based on RNA-seq signal in the bin (*n* = 576, *p*-values are from Tukey’s multiple comparisons test). Bins occupied by chromatin types 1–13 by less than 10% are not included in the analysis. RPK, reads per Kb; RPM, reads per Mb. (**F**) Distribution of raw contacts of MIR3687 along Chrs 18 and 19 and fold enrichment compared to background at a 50 Kb resolution. Gene distribution, RNA-seq signal (1 Kb bin), and replication timing profile for K562 as determined by Repli-seq (Hansen et al. 2010) are shown. (**G**) Distribution of correlation coefficients upon comparison of MIR3687 fold enrichment profile with Repli-seq in genomic windows of 20 Mb, as examined by StereoGene (Stavrovskaya et al. 2017). The genome-wide correlation coefficients calculated with the kernel and *p*-values are presented. (**H**) Fold enrichment of MIR3648, MIR3687, and U2 at individual chromosomes relative to overall contact frequency of respective RNAs in the genome. Error bars, SEM for two biological replicates.

### Cis and trans contacts of mRNAs

We next studied genomic contacts of exonic and intronic regions of mRNAs. In K562 cells, we identified 3.1M and 26.6M RNA–DNA chimeric molecules with RNA parts representing fragments of exons and introns, respectively. Occasionally, RNA parts intersected exon–exon or exon–intron junctions (∼0.8M RNA–DNA chimeras of each type) representing fragments of spliced or unspliced transcripts. We grouped RNA parts of chimeric molecules of each type according to their parental chromosome and determined how frequently their respective DNA parts are mapped to the same or other chromosomes. In a similar way, we analyzed Hi-C data for K562 cells (Rao et al. 2014). We identified DNA–DNA ligation products with one side mapped to exons or introns of the protein-coding genes lying on one chromosome and calculated how frequently the other side of the ligation product is mapped to the same or other chromosomes. In this way, we determined frequencies of cis and trans contacts for exon and intron regions of mRNAs and protein-coding genes (Table S6). The results were presented as averages for all chromosomes (Fig. 4A). It became clear that, although both mRNAs in Red-C data and protein-coding genes in Hi-C data show a clear preference for cis contacts, the former interact with non-parental chromosomes 10–20 times more frequently than the latter (Fig. 4A; see also Fig. S7). Accordingly, within their own chromosome, mRNAs interact with remote regions more frequently than their own genes, as follows from the analysis of scaling of contact probabilities showing a slower slope for mRNA curves (Fig. 4B). Hence, an mRNA does not occupy the gene most of the time and does not merely follow its interaction pattern; a significant portion of contacts occur when an mRNA is released from the gene. Remarkably, longer mRNAs are characterized by a higher proportion of cis to trans contacts, apparently due to a longer linkage with the parental chromosome in the course of transcription (Fig. 4C). Also of note is that exons of mRNA, especially those present in spliced transcripts, show a higher frequency of inter-chromosomal contacts than introns, especially those present in unspliced transcripts (Fig. 4A, Fig. S7). Finally, although the total number of contacts is higher for introns compared to exons (apparently due to higher intron length), exons establish ∼2 times more contacts than introns per RNA unit length (Fig. S8). These results likely reflect the different fate of exons, which are included into mature mRNA and occasionally contact multiple genomic sites during mRNA export from the nucleus, and introns, which are rapidly degraded.

**Fig. 4.**
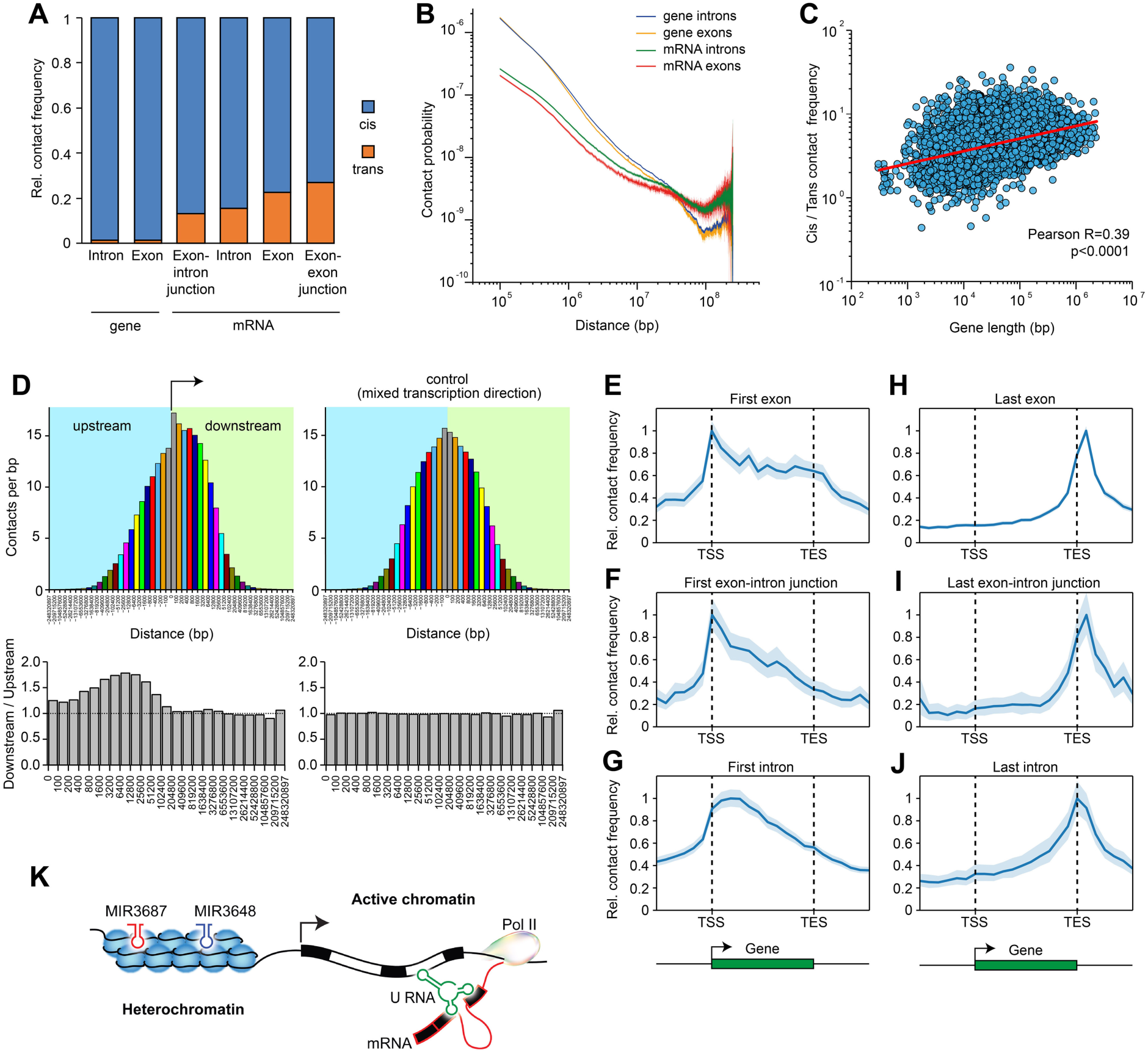
Inter- and intra-chromosomal contacts of mRNAs. (**A**) Relative frequency of cis and trans contacts for different regions of mRNAs and protein-coding genes averaged for all chromosomes. See also Fig. S7. (**B**) Double logarithmic scaling plot of the dependence of contact probability on genomic distance for exons and introns of mRNAs and protein-coding genes. Colored area in the background of curves, 95% CI. (**C**) Correlation between length of protein-coding genes and ratio between frequencies of cis and trans contacts for mRNAs encoded by these genes. (**D**) Frequency of contacts of mRNA fragments with downstream and upstream intervals with (left) or without (right) respect to the direction of transcription. Pairs of bars of the same color represent results for equally spaced regions downstream and upstream of mRNA fragments. Shown below are the ratios of contact frequencies between equally spaced regions downstream and upstream of mRNA fragments. (**E–J**) Contacts of the different regions of mRNA with the body of the encoding gene and its flanking regions averaged over all mRNAs establishing at least one contact with the gene body or flanking areas (*n* = 11,122). The maximal value of the averaged profile is taken to be equal to 1. Colored area in the background of curves, 95% CI. (**K**) Scheme illustrating the main findings of the study.

We next considered the contacts of mRNAs along the gene body. First, we determined frequencies with which mRNA fragments interact with genomic regions upstream and downstream of the site from where the fragment was transcribed, with respect to the direction of mRNA transcription. As expected, the highest interaction frequency was observed over the transcription site (Fig. 4D, upper left), which likely reflects the association of nascent RNA and DNA via transcription complex. The contact frequency decreases with an increase in distance from the transcription site, resulting in a characteristic bell-shaped distribution of contacts (Fig. 4D, upper left). Strikingly, the distribution is asymmetric relative to the transcription site. We found that the interaction frequency of mRNA fragments is ∼1.5 times higher in downstream regions as compared to upstream regions; the difference fades at distances more than 100 Kb from the transcription site (Fig. 4D, lower left). For a control, we calculated contact frequencies irrespective of transcription direction; the observed difference disappeared, and the distribution of contacts became symmetric relative to the transcription site (Fig. 4D, upper and lower right). These observations can be explained by the fact that RNA is dragged behind the RNA polymerase during transcription.

We further produced average profiles of mRNA binding over the body of encoding genes. With this aim, we divided protein-coding genes and their flanking regions into 24 bins and averaged the contacts of particular mRNA regions for bins located in the same position relative to the transcription start site (TSS) across all mRNAs. We started by examining the contacts of the first exon and intron with the downstream regions of the gene. The contact frequency of the first exon remains quite high until the transcription end site (TES) and sharply decreases beyond the TES (Fig. 4E). The first intron shows the same tendency; however, the contact frequency decreases more sharply within the gene body, and no break in the curve is distinguishable at the TES (Fig. 4G). The same is true for the first exon–intron junction (Fig. 4F). It thus appears that the first intron is co-transcriptionally removed from the transcript, while the first exon moves with the transcription complex up to the TES until the termination of transcription. In contrast, the last exon, last intron, and last exon–intron junction show almost the same decrease in contact frequencies toward the TSS, and in this case, the decrease within the gene body is as sharp as beyond the gene body (Fig. 4H–J). This observation seems to reflect a disengagement of mRNAs from the gene after the transcription of the last exon.

The conclusion about co-transcriptional intron splicing was confirmed when consecutive segments of mRNAs from the 5’ to 3’ end were examined (Fig. S9). Remarkably, exons show an increased interaction frequency with the region immediately downstream of the TES that is particularly prominent for the last exon (Fig. 4H, Fig. S9A). This observation may indicate that RNA polymerase II, which is known to continue transcription beyond the gene boundary, entrains mRNA before the latter is cleaved at a poly(A) signal and released.

Surprisingly, our data do not support a popular hypothesis about gene circularization aiming to facilitate transcription re-initiation (Hampsey et al. 2011). If looping between promoter and terminator occurred, the first exon and intron would demonstrate an increased frequency of interaction with the end of the gene, while the last exon and intron would demonstrate an increased frequency of interaction with the beginning of the gene, neither of which is seen in our data (Fig. 4E–J, Fig. S9A,C). The result holds true for both long and actively transcribed genes (Fig. S10).

### Comparison with fRIP data

To find the proteins that could be involved in RNA–DNA interactions, we examined the data of RNA immunoprecipitation experiments (fRIP) from (Hendrickson et al. 2016). This study provides data on RNA–protein interactions for 24 chromatin-associated and RNA-binding proteins in K562 cells. We found that most contacting RNAs identified with Red-C and RNAs establishing contacts with proteins in the fRIP experiment intersect (Fig. S11A, hypergeometric test *p*-value < 2.2e-16). We also observed that RNAs with the highest propensity to bind chromatin (defined as the ratio of contact number to RNA-seq signal) frequently interact with Polycomb proteins (SUZ12, EZH2), histone acetylase/deacetylase (CBP/HDAC1), and other proteins involved in the control of chromatin folding and dynamics (DNMT1, CBX3). A significant number of chromatin-associated RNAs have contacts with the RNA editing protein ADAR (Fig. S11B).

## Discussion

It is becoming increasingly evident that various non-coding RNAs play important roles in nuclear organization, chromatin architecture, and regulation of gene expression (Bergmann and Spector 2014; Bohmdorfer and Wierzbicki 2015; Jarroux et al. 2017; Sun et al. 2018; Cao et al. 2019). Still, it is likely that many functions of ncRNAs and many functionally significant individual ncRNAs are yet to be disclosed and characterized. The progress in this area of research depends on the availability of data on genomic/chromosomal distribution of various types of RNAs. Several studies aiming to characterize the RNA–DNA interactome have been published recently (Li et al. 2017; Sridhar et al. 2017; Bell et al. 2018). The experimental approach used in these studies is based on adapter-mediated proximity ligation of RNA to DNA within fixed nuclei. In all protocols published so far, only the 3’ end of captured RNA is identified. Here we present Red-C, a modified version of the adapter-mediated proximity RNA–DNA ligation protocol that allows for mapping of both the 5’ and 3’ ends of captured RNA fragments. This allows for identification of intermediate splicing products, products of alternative splicing and trans-splicing, and polyadenylated transcripts, as well as for discrimination of micro-RNAs from their precursors. The polarity of the Red-C procedure rules out the possibility of DNA–DNA and RNA–RNA ligation and unambiguously defines the position of RNA and DNA parts in the RNA–DNA chimeras. Red-C can be readily upgraded for selective enrichment of RNA–DNA libraries by the C-TALE protocol recently developed in our laboratory (Golov et al. 2019), thus providing opportunities for obtaining high-resolution contact profiles for any RNA(s) of interest.

Using the Red-C procedure, we identified a number of presently unknown sense and antisense RNAs interacting with DNA in the vicinity of structural genes as well as ncRNAs preferentially associated with specific chromatin types. The entirety of the data obtained is yet to be fully explored. Here, we began this analysis by partitioning chromatin-bound RNAs into groups according to their genomic distribution relative to their parental transcription unit. This kind of analysis (Fig. 2, Fig. S6) allowed us to distinguish potential regulatory RNAs acting locally from those acting genome wide. Indeed, the known trans- and cis-acting ncRNAs (such as sn and sno RNAs on one side and XIST on the other side) fell into distinct groups. Of particular interest could be the 313 RNAs enriched in gene-proximal areas. This group is enriched with vlinc RNAs and unannotated antisense X RNAs. Most of them show a preference for association with inactive chromatin regions and thus might be involved in silencing of the transcription of nearby genes. In the group of RNAs interacting with chromatin genome wide (224 ncRNAs), MIR3648 and MIR3687—or rather, longer precursors of these miRNA—drew our attention because they demonstrated a preference for interaction with repressed genomic regions (inactive late-replicating chromatin compartment) and, to a lesser extent, with Polycomb-repressed facultative heterochromatin. With the exception of pericentromeric heterochromatin and facultative heterochromatin repressed by Polycomb, the nature of mechanisms mediating repression of a significant part of the genome within lamina-associated domains (LADs) remains obscure (van Steensel and Belmont 2017; Leemans et al. 2019). The two ncRNAs identified in our analysis appear to be good candidates for global regulators of this repression (Fig. 4K). Previous studies have revealed multiple examples of miRNA involvement in transcriptional silencing of individual genes (Roberts 2014). However, MIR3648 and MIR3687 represent the first examples of miRNA associated with inactive chromatin genome wide. Interestingly, these ncRNAs originate from the 5’ external transcribed spacer of the 45S rRNA operon. This may provide a new key for the proposed role of the nucleolus in the assembly of repressed chromatin domains (Guetg and Santoro 2012).

An interesting observation made in our study is that eRNA transcribed from strong enhancers interacts with other strong enhancers, but not with promoters. This may signify that enhancers are assembled in spatial clusters even when they do not interact with promoters or interact with different promoters transiently. Another option is that eRNA is involved in establishing communication between enhancers. This supposition certainly deserves further investigation.

The results of our analysis allowed the tracing of the dynamics of structural gene transcription and splicing for the first time (Fig. 4K). Our results strongly support the model of co-transcriptional splicing (Bentley 2014) and thus call into question the possibility that pre-mRNAs may execute some regulatory function before being spliced (Scherrer 2018). This may not apply to circular RNAs (Li et al. 2018) that were not specifically analyzed in our study. Finally, our results do not support the model of gene circularization (Hampsey et al. 2011). Although we cannot exclude a possibility that in some specific cases the genes may be circularized, the majority of structural genes appear to remain linear in the course of transcription.

## Methods

### Red-C procedure

Approximately 2.5×10^6^ cells were cross-linked with 1% formaldehyde (Sigma-Aldrich F8775) in full growth media for 10 min at room temperature followed by quenching with 125 mM glycine. Cells were washed with cold PBS and incubated in 375 µL lysis buffer (10 mM Tris pH 7.5, 10 mM NaCl, 0.2% NP40, 1× protease inhibitors (Bimake), 37.5 U SUPERase.In RNase inhibitor (Invitrogene)) for 10 min on ice. To remove cytoplasm and extract RNA and proteins that were not cross-linked to DNA, permeabilized cells were resuspended in 250 µL nuclease-free water (Qiagen) followed by adding 7.5 µL 10% SDS and incubated for 30 min at 37 °C with shaking at 1200 rpm. SDS was sequestered by adding 25 µL 20% Triton X-100 followed by incubation for 30 min at 37 °C with shaking at 1200 rpm. After adding 100 µL warm 4× NEB buffer 4, nuclei were pelleted for 3 min at 2500 g and resuspended in 250 µL 1× NEB buffer 4. DNA was digested by adding 10 µL NlaIII (10 U/µL, NEB) and incubated for 3½ h at 37 °C with shaking at 1200 rpm. Nuclei were pelleted as above and resuspended in 150 µL 1× NEB buffer 2 followed by adding 3.75 µL 10% SDS to inactivate residual restriction enzyme.

Nuclei were immobilized on magnetic beads by mixing the suspension with 310 µL AMPure XP beads (Beckman Coulter) and incubating for 5 min at room temperature. Immobilization on beads helps to manipulate nuclei in downstream steps (Ramani et al. 2016). It does not influence the procedure’s performance because carrying out the experiment without the beads produced the same results (data not shown). Bead-nuclei were collected on a magnet, washed twice with 1 mL 80% ethanol and, after removing residual ethanol by 10 s spinning at 500 g, air-dried for 1 min. 3’ P ends of RNA were dephosphorylated by resuspending bead-nuclei in 190 µL dephosphorylation solution (1× PNK buffer (NEB), 0.1% Triton X-100, hereinafter the concentration is given as for the enzyme-containing mixture), followed by adding 10 µL PNK (10 U/µL, NEB). The mixture was incubated for 30 min at 37 °C with shaking at 800 rpm. Bead-nuclei were pelleted for 2 min at 2500 g and resuspended in 189 µL blunting solution (1× T4 DNA ligase buffer (NEB), 0.25 mM dNTPs, 0.1% Triton X-100). The mixture was supplemented with 5 µL DNA polymerase (3 U/µL, NEB) and 6 µL Klenow (5 U/µL, NEB), and DNA blunting was carried out for 1 h at room temperature with shaking at 800 rpm. The reaction was stopped by adding 5 µL 10% SDS followed by pelleting bead-nuclei as above. Bead-nuclei were washed with 200 µL 1× NEB buffer 2 supplemented with 1% Triton X-100, pelleted, and resuspended in 198 µL A-tailing solution (1× NEB buffer 2, 500 mM dATP, 1% Triton X-100). DNA ends were A-tailed by adding 1.5 µL Klenow (exo-) (50 U/µL, NEB) followed by incubation for 1 h at 37 °C with shaking at 800 rpm. Bead-nuclei were subsequently washed with 200 µL 1× RNA ligase buffer (NEB) supplemented with 0.1% Triton X-100 and with 200 µL 1× RNA ligase buffer (NEB) by repeating resuspending/pelleting.

The 3’ OH ends of RNA were ligated with 5’ rApp ends of a bridge adapter (a duplex of 5’-/rApp/TCCTAGCACCATCAATGCGATAGGCAACGCTCCGACT-3’, 3’ hydroxyl non-blocked, and 5’-/Phos/GTCGGAGCGTTGCC/T-Biotin/ATCG-3’). For this purpose, bead-nuclei were resuspended in 190 µL RNA ligase solution (1× RNA ligase buffer (NEB), 4.5 µM bridge adapter, 20% PEG-8000 (NEB)), 10 µL T4 RNA ligase 2 truncated (200 U/µL, NEB) was added, and the mixture was incubated for 6 h at room temperature then overnight at 16 °C with shaking at 800 rpm. To wash off non-ligated bridge adapter, bead-nuclei were pelleted, resuspended in 200 µL nuclease-free water and mixed with 165 µL AMPure buffer (20% PEG-8000, 2.5 M NaCl) (Ramani et al. 2016). Bead-nuclei were collected on a magnet, washed once with 1 mL 80% ethanol, resuspended in 200 µL nuclease-free water and again mixed with 165 µL AMPure buffer. Bead-nuclei were collected on a magnet, washed twice with 1 mL 80% ethanol, and resuspended in 95 µL PNK solution (1× T4 DNA ligase buffer (NEB), 0.1% Triton X-100). Then, 5 µL PNK (10 U/µL, NEB) was added, and the mixture was incubated for 1h at 37 °C with shaking at 800 rpm. Bead-nuclei were pelleted, resuspended in 980 µL 1.02× T4 DNA ligase buffer (Thermo Scientific), and split into 2 equal portions. To one portion 10 µL T4 DNA ligase (5 Weiss U/µL, Thermo Scientific) was added, to the other 10 µL nuclease-free water (DNA ligase minus control). DNA proximity ligation was allowed to proceed for 6 h at room temperature with rotating, followed by pelleting bead-nuclei for 5 min at 7400 g.

To reverse formaldehyde cross-links and digest proteins, bead-nuclei were resuspended in 235 µL proteinase K solution (100 mM NaCl, 10 mM Tris pH 7.5, 2 mM EDTA, 1% SDS), 15 µL proteinase K (20 mg/mL, Ambion) was added, and incubation for 1 h at 55°C and then for 2 h at 65 °C followed. To precipitate RNA–DNA chimeras, 1.5 μL GlycoBlue (Thermo Scientific), 25 μL 3M NaAC and 275 μL isopropanol were added and, after overnight incubation at −80 °C, the mixture was centrifuged for 30 min at 16100 g and 4 °C. The pellet was resuspended in 50 μL nuclease-free water, and RNA–DNA chimeras were further purified with 2 volumes of AMPure XP beads and finally eluted into 50 μL nuclease-free water followed by measuring the concentration with a Qubit dsDNA broad range kit. For the control experiment with RNase A treatment, RNA–DNA chimeras (3.5 μg) were incubated with 0.4 μL RNase A (10 mg/ml, Thermo Scientific) in water for 30 min at 37 °C, followed by clean up with 2 volumes of AMPure XP beads.

RNA–DNA chimeras (3.5 μg) were digested with MmeI in 100 μL reaction containing 1× NEB buffer 4, 0.2 mg/mL BSA (NEB), 80 μM SAM (NEB), 0.1 μM ds oligo with MmeI site (a duplex of 5’-CTGTCCGTTCCGACTACCCTCCCGAC-3’ and 5’-GTCGGGAGGGTAGTCGGAACGGACAG-3’), and 4 U MmeI (NEB) for 2 h at 37 °C. Short dsDNA containing the MmeI site is added to stimulate the cleavage of DNA molecules containing a single MmeI site (Morgan et al. 2008).

After MmeI digestion, RNA–DNA chimeras were subjected to biotin pull-down. For this process, 10 μL of Dynabeads MyOne Streptavidin C1 beads (10 mg/mL, Thermo Scientific) was washed twice with 400 μL tween washing buffer (TWB) (5 mM Tris pH 7.5, 0.5 mM EDTA, 1 M NaCl, 0.05% Tween 20) by repeating the resuspension/magnet separation. Streptavidin beads were resuspended in 100 μL 2× binding buffer (10 mM Tris pH 7.5, 1 mM EDTA, 2 M NaCl) and mixed with the solution after MmeI digestion followed by incubation for 15 min at room temperature to bindbiotinylated bridge to streptavidin beads. Streptavidin beads with tethered RNA–DNA chimeras were washed twice with 600 μL TWB, once with 100 μL 1× NEB buffer 2, once with 50 μL 1× First-Strand Buffer (Clontech) and resuspended in 38 μL reverse transcriptase solution (1× First-Strand Buffer (Clontech), 2.5 mM DTT (Clontech), 1 mM dNTPs, 1 μM switch template oligo (5’-iCiGiCGTGACTGGAGTTCAGACGTGTGCTCTTCCGATCTrGrGrG-3’ where iC and iG designate Iso-dC and Iso-dG, and r indicates ribonucleotides), 20 U SUPERase-In RNase inhibitor (Invitrogene)). After pre-heating at 42 °C for 2 min, reverse transcription was initiated from the bridge 3’ OH by adding 2 μL SMARTScribe Reverse Transcriptase (100 U/ μL, Clontech) and incubating for 1 h at 42 °C with shaking at 800 rpm. Reverse transcriptase first transcribes bridge DNA, then DNA-RNA junction, then RNA. Upon reaching the 5’ end of the RNA, reverse transcriptase adds a few non-template nucleotides (predominantly dC) to the 3’ end of cDNA. This dC stretch pairs with rGrGrG of the switch template oligo, and reverse transcriptase continues replication using the switch template oligo as a template (Zhu et al. 2001) (Fig. S1A). Atypical nucleotides isocytidine and isoguanine prevent secondary switching at the 5’ end of the switch template oligo (Kapteyn et al. 2010).

After cDNA synthesis, streptavidin beads were washed twice with 600 μL TWB, once with 100 μL 1× NEB buffer 2, once with 100 μL 1× T4 DNA ligase buffer (Thermo Scientific) and resuspended in 48 μL DNA ligase solution (1× rapid ligation buffer (Thermo Scientific), 3 μM NN-adapter (a duplex of 5’-AGATCGGAAGAGCGTCGTGTAGGGAAAGAGTGTAGATCTCGGTGGTCGCCGTATC ATT-3’ and 5’-AATGATACGGCGACCACCGAGATCTACACTCTTTCCCTACACGACGCTCTTCCGAT CTNN-3’ where N designates any base). To ligate DNA NN ends produced by MmeI digestion to adapter NN ends, 2 µL T4 DNA ligase (5 Weiss U/µL, Thermo Scientific) was added followed by incubation for 1 h at room temperature. The NN-adaptor is used in a non-phosphorylated form to avoid adaptor-to-adaptor ligation. As a result, a nick is left in the non-biotinylated strand (see Fig. S1A). After ligation, streptavidin beads were washed twice with 600 μL TWB, once with 100 μL 1× NEB buffer 2, once with 100 μL 10 mM Tris pH 8.0 and resuspended in 20 μL 10 mM Tris pH 8.0.

DNA-cDNA chimeras were amplified in 50 μL PCR containing 1× KAPA HiFi Fidelity Buffer, 0.3 μM dNTPs, 0.5 μM universal primer (5’-AATGATACGGCGACCACCGAGATCTACACTCTTTCCCTACACGA-3’), 0.5 μM indexed primer (5’-CAAGCAGAAGACGGCATACGAGATNNNNNNGTGACTGGAGTTCAGACGTGTGC-3’ where NNNNNN is a sequencing index), 1 U KAPA HiFi DNA Polymerase, and 4 μL streptavidin beads from the above step. PCR was performed as follows: 95 °C 5 min, 12–14 cycles of 98 °C 20 s, 65 °C 15 s, 72 °C 45 s, 72 °C 3 min. PCR products of 4 reactions were pooled, purified with 1.8 volumes of AMPure XP beads and separated in a 2% agarose gel. PCR products were excised from the gel and purified using a QIAquick Gel Extraction Kit (Qiagen). Purified PCR products were paired-end sequenced on the HiSeq 2500 or MiSeq Illumina platform with the read length of 80–133 n.

In the case of fibroblasts, we used 0.3 μg RNA–DNA chimeras for MmeI digestion and 17 cycles of PCR for final amplification.

All oligos were purchased from Integrated DNA Technologies, Inc. Certified RNAse-free reagents and materials were used.

### Read filtering and mapping

For filtering out possible PCR duplicates, both forward (R1) and reverse (R2) reads were cut to the first 50 nucleotides. Then, fastuniq was used for searching for exact duplicates. From a group of duplicates, only one pair was retained.

For sequencing quality control, we ran FASTQC and found a decline in sequencing quality at the end of reads. We used TRIMMOMATIC for the detection of the first leftmost low-quality position in each forward and reverse read. The parameters were set to: window size: 5, quality threshold: 26. Only reads with at least one nucleotide passing the quality control filter were selected.

Each read, regardless of quality filter, was subjected to the scanning of adaptors, bridge, and GGG/CCC oligonucleotides. The scanning was done with Rabin–Karp algorithm implementation in C. For that, sequences from FASTQ files with reads and FASTA files with oligonucleotides were first converted to binary indexed files; then, the search was run. For each read, the positions of start and end of oligonucleotides were reported. First, R1 was scanned for a complete bridge (AGTCGGAGCGTTGCCTATCGCATTGATGGTGCTAGGA). The bridge was allowed to have 1 mismatch in any position except the rightmost GA and to be positioned anywhere in R1. If a complete bridge was identified in R1, the read pair was retained. Second, only read pairs with an R2 read starting with GGG were retained. Third, R1 was scanned for STO (CCCAGATCGGAAGA was required) with allowing 1 mismatch. If identified, R1 was trimmed right in front of STO. To trim shorter pieces of STO that could occur at the end of R1, we took 14 nucleotides adjoining GGG in the start of R2, converted to reverse complement, and performed scanning of R1 for the rightmost position of complementarity. If identified, the region of R1 to the right of the position of complementarity was trimmed. Fourth, R2 was scanned for the bridge (TCCTAGCACCATCA was required) with allowing 1 mismatch. If identified, R2 was trimmed right in front of the bridge. To trim shorter pieces of a bridge that could happen at the end of R2, we took 14 nucleotides located to the right of the bridge in R1, converted to reverse complement, and performed scanning of R2 for the rightmost position of complementarity. If identified, the region of R2 to the right of the position of complementarity was trimmed. Finally, we extracted the DNA part as the region of R1 to the left of the bridge, the RNA 3’ part as the region of R1 to the right of the bridge, and the RNA 5’ part as the region of R2 to the right of the first GGG. Note that RNA 3’ and RNA 5’ parts can partly or completely overlap if the RNA portion of the chimera is short. If the lengths of DNA, RNA 3’, and RNA 5’ parts were more than 0, these sequences were written to separate FASTQ files with corresponding qualities from initial files.

Most DNA parts are 18–20 nucleotides long (Fig. S1E), with the length distribution precisely following the MmeI digestion pattern (Fig. S1F, left graph). Indeed, MmeI cuts predominantly at 20 and 21 bp upstream of the recognition site with approximately a 60%/40% ratio; there is also minor cutting at 19 bp (a few percent). A 1 bp shift (18–20 instead of 19–21) is consistent with the position of the MmeI site in the bridge. DNA parts of 0 or 1 nucleotide are also observed (Fig. S1F, left graph). They represent chimeras without the DNA part and can be seen in the control experiment without DNA ligase (Fig. S1C). In contrast to DNA parts, RNA parts demonstrate a wide range of length distributions (Fig. S1F, right graph). Short RNA parts may represent short RNA species or result from fragmentation of RNA during multiple incubations at 37 °C in the presence of Mg++, likely by a chemical mechanism. Of note, performing all steps of the experimental procedure in the presence of SUPERase-In RNase inhibitor did not increase the average size of RNA parts (data not shown). More importantly, in the control experiment with RNase A, in the majority of reads, RNA parts were absent or did not exceed a few nucleotides (Fig. S1E), which are always A and/or G (manifested by T/C in the forward read, Fig. S1C). This result was expected from the RNase A digestion mechanism; RNase A cleaves RNA at pyrimidines, thus leaving purine ribonucleotides adjacent to the bridge preserved. Overall, the above observations argue for the validity and specificity of the developed protocol.

DNA parts of 18–20 nucleotides, RNA 3’ parts of ≥14 nucleotides, and RNA 5’ parts of ≥14 nucleotides were independently mapped to the hg19 genome with the hisat2 program. Before mapping, the end of the DNA part adjoining the bridge was supplemented with CATG (the 3’ overhang produced by NlaIII digest and then blunted, see Fig. S1A) to increase the yield of unique mappings. We used parameters: “-k 100 --no-spliced-alignment --no-softclip” for DNA and “-k 100 --no-softclip --dta-cufflinks” for RNAs (--known-splicesite-infile; the splicing site annotation was taken from gencode v19). SAM files were filtered for unique mappings with at most 2 mismatches relative to the reference genome with samtools and converted to BED with bedtools. We retained only such DNA-RNA 3’-RNA 5’ triples that were all successfully and uniquely mapped to the canonical chromosomes. If one of the parts was missing, non-uniquely mapped, unmapped, or mapped to the non-canonical chromosome, the read pair was filtered out (Fig. S1G, Table S1).

We found that the end of the RNA 3’ part adjoining the bridge and the end of the RNA 5’ part adjoining GGG, which mark correspondingly the 3’ and 5’ ends of RNA in the chimera, are a slightly more frequently mapped to NlaIII sites than may be expected based on random distribution (Fig. S1H), an observation that may be indicative of the traces of DNA-DNA ligation in the procedure. We thus discarded a read pair if the 3’ or 5’ or both ends of the RNA fell within the NlaIII site ± 1 letter. We also discarded a read pair if the 5’ end of the RNA fell within the MmeI digestion site because that MmeI site may naturally occur within a NlaIII fragment. It is obvious that concomitantly, we lost a fraction of genuine RNA–DNA ligation products because an RNA end may occasionally happen within NlaIII and MmeI digestion sites. However, at that cost, we eliminated possible experimental artifacts. Finally, to avoid spurious trans-contacts that could originate from intermolecular template switching of the reverse transcriptase (Houseley and Tollervey 2010), we required that RNA 3’ and RNA 5’ parts be mapped to the opposite strands of the same chromosome at a distance of no more than 10 Kb from each other (as measured by the difference between the lower coordinates of mapping). For K562 cells, we retained a read pair if the DNA part mapped to genomic regions annotated by chromatin states 1–13 (Ernst et al. 2011). If the DNA part mapped to repetitive/CNV chromatin (chromatin states 14-15) or beyond annotated regions, the read pair was filtered out (<1.5% of all read pairs).

Table S1 shows the number of read pairs retained after each consecutive step of the data processing pipeline described above.

### Annotation of RNAs

We use RNA 3’ parts retrieved from the forward reads as described above. If a splicing junction is reported within the RNA 3’ part, we use a fragment of the RNA 3’ part from the bridge to the break in alignment. We intersect RNA 3’ parts with the following gene tracks: gene annotation from GENCODE (release 27 (GRCh37); basic gene annotation); piRNAs annotation from piRNABank; tRNAs annotation from UCSC (track: tRNA Genes; table: tRNAs); set of rRNAs, snRNAs, scRNAs, tRNAs, RNAs, srpRNAs from UCSC (track: RepeatMasker; table: rmsk); vlinc from Laurent et al. (St Laurent et al. 2013). In case the RNA 3’ part intersects a gene by at least 1 nucleotide, this RNA part is assigned to this gene (we require that the RNA 3’ part be mapped to the strand opposite to that of the gene as expected from the Red-C procedure, see Fig. S1A). If the RNA 3’ part intersects more than one gene at the same strand, this RNA 3’ part is assigned to the gene showing the highest coverage by RNA parts, which is determined as the total number of RNA 3’ parts mapped to the gene normalized to the gene length. In this way, we ensure that RNA parts representing highly expressed small RNAs (such as U snRNAs) are not assigned to the genes hosting these small RNA genes. At the final step, we combine DNA parts mated with RNA 3’ parts originating from a single gene, thus obtaining a whole-genome contact profile for each respective RNA.

Clusters of RNA parts that were not assigned to any gene may potentially represent novel chromatin-associated RNAs (designated X-RNAs). To identify X-RNAs, we search for clusters comprising at least 100 non-assigned RNA 3’ parts mapped to the same strand, with a distance between consecutive RNA parts of no more than 100 bp. If a known gene is detected at a distance of less than 100 bp of the cluster boundaries that is covered by more RNA parts than there are in the cluster, the cluster is discarded because it may represent a “tail” of this gene. Clusters spaced less than 1 Kb apart at the same strand are further aggregated into one cluster to compensate for coverage gaps. The procedure yields 1867 X-RNAs in K562 cells (Table S3).

eRNAs are arbitrarily defined as RNAs produced from an enhancer-specific chromatin type (states 4-7) (Ernst et al. 2011). Each genomic interval annotated by chromatin states 4, 5, 6, or 7 is considered an individual enhancer (medium length 1400, 800, 600, and 1400 bp, respectively). RNA 3’ parts mapped to either strand of so-defined enhancers are assigned to these enhancers independently of whether they are assigned to any other gene. If the RNA 3’ part intersects an enhancer and some gene, we count this RNA 3’ part twice as a part of the eRNA and a part of the RNA encoded by this gene. We identify 9063 eRNAs with ≥ 100 contacts (Table S4).

### Construction of background, normalization, and enrichment calculation

To account for the level of background ligation in the procedure, we estimate the total number of mRNA trans-contacts with each genomic site, as suggested by Li et al. (Li et al. 2017). We divide the genome into 500 bp bins, and for each bin, we sum the number of contacts made with this bin by protein-coding RNAs originating from all over the genome except the chromosome where the bin belongs. We smooth the obtained signal with a Gaussian function and use it as a background signal. We then normalize raw counts of individual RNA–DNA contacts by the value of the background in the genomic coordinate where the DNA part is mapped. To work with DNA parts mapped to regions with zero value of the background signal (< 0.01% of all DNA parts), we add to the denominator a pseudocount constituting ∼ 10% of the minimal non-zero value of the background (∼ 0.0001% of the mean value of the background). Finally, we divide the sum of raw counts by the sum of normalized counts and multiply each normalized count by the obtained coefficient. In such a manner, the sum of normalized counts is equalized with the sum of raw counts, whereas each contact of the library is rescaled according to the background level.

To determine the average contact frequency of a given RNA in a region of interest (e.g., gene, parental chromosome, the full genome), we sum the number of background normalized contacts this RNA establishes with the region and divide by the total length of the region. In our analysis, we discard contacts with genomic regions annotated by chromatin types 14 and 15 and not annotated by any chromatin type. If such regions occur within the region of interest, we subtract the length of these regions from the total length of the region.

To calculate the enrichment of an individual RNA compared to the background, we use the procedure described by Li et al. (Li et al. 2017) with minor modifications. We divide the genome into bins of an appropriate size, and for each bin, we sum the number of contacts made with this bin by protein-coding RNAs originating from all over the genome except the chromosome where the bin belongs. We normalize the signal in each bin by the average value of the signal among all bins. We smooth the obtained signal with a moving window of 10 bins and use it as a background signal. We next calculate the number of contacts of a selected RNA with each bin and normalize by the average number of contacts of this RNA among all bins. We then divide the signal for the selected RNA by the signal for background in each bin, thus yielding the fold enrichment of this RNA compared to the background. To work with robust enrichment, we filter out bins with fold enrichment < 2. We further retain bins meeting the following requirement: at least 3 bins with fold enrichment ≥2 in the 11-bin window centered on the bin. Finally, we smooth the signal by a sliding window of 10 bins. In this way, we identify peaks of enrichment of individual RNAs along the genome.

### Chromatin types

We use the annotation of chromatin states for K562 cells obtained by Ernst et al. (Ernst et al. 2011). The authors of that study used combinations of chromatin marks to divide the genome into 15 non-overlapping chromatin states: active promoters (1), weak promoters (2), inactive/poised promoters (3), strong enhancers (4 and 5), weak enhancers (6 and 7), CTCF-dependent insulators (8), transcriptional transition (9), transcriptional elongation (10), weak transcribed (11), Polycomb repressed (12), bulk heterochromatin (13), and repetitive/CNV (14 and 15). We consider individual chromatin types from 1 to 13 and their combinations: 1+2+4+5+6+7+9+10+11 for active chromatin, 3+12 for Polycomb repressed chromatin, and 3+12+13 for repressed chromatin.

To determine the average contact frequency of an RNA with a particular chromatin type, we sum the number of background normalized contacts with this chromatin type in a region of interest and divide by the total length of this chromatin type in the region of interest.

### Ranking of RNAs by preference for short- and long-range contacts

For each RNA, we consider several genomic intervals: the region encoding for this RNA (gene, G); 0–500 Kb upstream and downstream of gene boundaries (short & medium cis, SM); 500 Kb–5 Mb upstream and downstream of gene boundaries (long cis, L); > 5 Mb from gene boundaries in the same chromosome (remote cis, R); and finally the other chromosomes (trans, T) (Fig. 2A).

We select RNAs with ≥ 500 contacts in total and at least 1 contact in each of the following intervals: L, R, and T (10367 RNAs, listed in Table S5). For each RNA, we calculate the average contact frequency in intervals SM, L, R and T, followed by computation of the ratios SM/L, L/R, and R/T. Considering the incline of point clouds in Fig. 2B, we divide RNAs into 3 groups with a low (500–1500), medium (1500–10000) and high (>10000) number of contacts. We Z transform SM/L ratios within each group, divide the obtained values into 5 quantiles, combine RNAs belonging to the same quantile for the 3 groups, and finally rank RNAs according to their SM/L value. We repeat the procedure for L/R and R/T ratios. An RNA is considered enriched in gene proximal area (± 5 Мb from gene) if it falls into the first quantile by SM/L value and into the fifth quantile by L/R and R/T values (group A, Fig. S6A). An RNA is considered XIST-like if it falls into the first quantile by SM/L and L/R values and into the fifth quantile by R/T value (group B, Fig. S6B). An RNA is considered distributed throughout the genome if it falls into the first quantile by each of the three values (group C, Fig. S6C).

### Contacts of exons and introns of mRNA

We distinguish 4 classes of RNA parts based on the mapping position of its 5’ end (the end adjoining GGG) and 3’ end (the ends adjoining bridge): (i) both 5’ and 3’ ends are within the same exon; (ii) both 5’ and 3’ ends are within the same intron; (iii) the 5’ end is within an exon and the 3’ end is within the next intron or the 5’ end is within an intron and the 3’ end is within the next exon (exon-intron junction); and (iv) the 5’ end is within an exon and the 3’ end is within the next exon, with a splicing junction reported within the RNA part (exon-exon junction). We also discriminate RNA parts representing different portions of mRNA, such as the first or last exon/intron, or a particular bin (RNA parts are assigned to a bin based on the position of the RNA 3’ end).

In the analysis of intra-gene contacts, we select protein-coding mRNAs that establish at least 1 contact with its own gene or gene flanking regions of half gene length (if there are several isoforms, we use the longest according to RefSeq annotation). We divide corresponding genomic regions into 24 bins (12 bins for gene body ± 6 bins for flanks). Note that bin length varies depending on gene length. For each mRNA, we consider RNA parts of a given type (e.g., pieces the first exon, intron regions of the second bin, etc) and determine the number of background normalized contacts these RNA parts establish with each genomic bin. Finally, we average contacts for genomic bins located in the same position relative to the gene body for all mRNAs.

### Scaling of contact probabilities

For calculating the scaling of contact probability of exons of mRNAs with regions of the chromosome bearing the encoding gene, we select a set of RNA–DNA contacts such that the 3’ end of the RNA part is mapped within an exon of a protein-coding gene and the DNA part is mapped anywhere in the genome. We divide the genome into 100 Kb bins and select bins to which one or more RNA parts are mapped. We then consider DNA parts mated to RNA parts of a given bin and calculate how many of these DNA parts are mapped to each consecutive genomic bin (zero values are recorded as well), thus yielding the number of contacts RNA of a given bin establishes throughout the genome. Finally, the contact numbers are averaged among pairs of bins equally spaced in a linear chromosome, and obtained values are normalized to the total number of contacts in the set, including both cis and trans-contacts. Note that the number of bin pairs decreases with an increase in distance between bins because short distances are assessed from both short and long chromosomes, whereas long distances are assessed only from long chromosomes.

Scaling of contact probabilities for introns of mRNAs is calculated in the same way with the only difference being that we select a set of RNA–DNA contacts such that the 3’ end of the RNA part is mapped within an intron of a protein-coding gene. To calculate the scaling of DNA-DNA contacts, we used a publicly available Hi-C data set for K562 (Rao et al. 2014). We require that one part of the DNA-DNA ligation product be mapped to an exon/intron of a protein-coding gene and the other part of DNA-DNA ligation product be mapped anywhere in the genome. Other steps of the analysis are done as described above for RNA–DNA contacts.

### Cell culture

Human K562 cells (ATCC^®^ CCL-243™) were cultured in RPMI supplemented with 10% FBS and penicillin/streptomycin. Normal human skin fibroblasts (female 46XX) were kindly provided by Dr M. Lagarkova (Federal Research and Clinical Center of Physical-Chemical Medicine, Moscow, Russia) and were cultured in DMEM supplemented with 10% FBS and penicillin/streptomycin. Human cells were grown at 37 °C and 5% CO_2_ in a conventional humidified CO_2_ incubator. Drosophila melanogaster Schneider-2 (S2) cells were a kind gift of Dr O. Maksimenko (Institute of Gene Biology, Moscow, Russia) and were grown at 25 °C in Schneider’s Drosophila Medium supplemented with 10% FBS and penicillin/streptomycin.

### RNA-seq

Total cellular RNA was isolated from K562 cells using an RNeasy Mini Kit (Qiagen) followed by the removal of ribosomal RNA using a Ribo-Zero Gold rRNA Removal Kit (Human/Mouse/Rat) (Illumina). Strand-specific sequencing libraries were prepared using a NEBNext Ultra II Directional RNA Library Prep Kit for Illumina (NEB). Libraries from two biological replicates were sequenced on the Illumina NextSeq 500 platform resulting in 22-25×10^6^ single-end reads. Reads were mapped and annotated by genes in the same way as for the RNA 3’ parts of the RNA–DNA chimeras (see above).

## Data availability

Datasets with raw fastq Red-C and RNA-Seq data and processed TSV files with contacts are available under GEO accession: GSE136141.

The code for read processing is available as RedClib on github: https://github.com/agalitsyna/RedClib

## Acknowledgements

This study was supported by the Russian Science Foundation (grant 18-14-00011). AAGal was supported by the Skoltech Center of Life Sciences Systems Biology Fellowship Program. This study was performed using the equipment of the Center for Precision Genome Editing and Genetic Technologies for Biomedicine of the Institute of Gene Biology RAS. The authors would like to acknowledge the computational resource Makarich provided by the Faculty of Bioengineering and Bioinformatics of Lomonosov Moscow State University and its administrators.

## Author contribution

AAM, SVR, and AAGav conceived of the study; AAGav developed the Red-C protocol; AAGal processed sequencing data; AAZ, AAGal, AVL, NMR, and MDM performed bioinformatics analysis under the supervision of AAM, AAGav, OLK, and SVU; MDL performed RNA-seq and NGS; NVP and AKG carried out cell culture work; AAGav and SVR wrote the manuscript with input from all authors.

